# Cell-state specific drug-responses are associated with differences in signaling network wiring

**DOI:** 10.1101/2025.01.27.635060

**Authors:** Niels Krämer, Roderick van Eijl, Tim Stohn, Sabine Tanis, Lodewyk Wessels, Evert Bosdriesz, Klaas W. Mulder

**Affiliations:** Radboud Institute for Molecular Life Sciences, Faculty of Science, Department of Molecular Developmental Biology, Radboud University, Nijmegen, the Netherlands; Department of Computer Science, Vrije Universiteit, Amsterdam, the Netherlands; Department of Molecular Carcinogenesis, Netherlands Cancer Institute, Amsterdam, the Netherlands

## Abstract

Intracellular signaling pathways form networks through which information is transmitted, often in the form of kinase-mediated phosphorylation events, to interpret extracellular signals and elicit appropriate cellular responses. Yet, even isogenic cells in a homogenous environment show heterogeneity in their intracellular “cell-state”, as well as their response to extracellular signals. Here, we aimed to better understand this relation between these phenomena by investigating how information flows through the EGF-receptor centered network upon targeted drug treatment, and how this is affected by cell-to-cell-state differences. Using single-cell ID-seq, we profiled the cell-state and signaling activity of primary human epidermal stem cells by measuring 69 (phospho-)proteins upon inhibition of the Erk/MAPK (p90RSK) and Akt/mTOR (p70S6K) routes downstream of the EGF pathway. We found that the effects of drug treatment propagated from the EGF-signaling pathway to other connected parts of the cellular signaling network, indicating altered signaling flow. We identified nine distinct cell-states that show pervasive state-dependent drug-responses for many (phospho-)proteins. Computational modeling of the signaling network using single-cell Comparative Network Reconstruction showed that many interactions between phospho-proteins (i.e. network wiring) were quantitatively different between cell-states. Furthermore, (phospho-)proteins with a cell-state dependent drug response, were more likely to be involved in interactions that showed a cell-state dependent strength. Overall, our results indicate that drug treatment response and signaling interactions between proteins are closely related and modulated by cell-state.

## Introduction

Cells in the human body constantly receive signals from their environment and have to translate these cues into correct cellular responses.What determines how a cell will react to a signal is still poorly understood. Many of the major communication signals that cells use are recurrent throughout embryonic development, in normal physiology, and are frequently mis-regulated in disease. In each of these biological contexts, these signals can have very different, even opposing, consequences. Cellular responses start with interpretation of environmental signals. In many cases, extracellular peptide growth factors bind to a cell’s surface receptors, initiating a cascade of phosphorylation events that eventually converge on transcription factors in the nucleus, leading to changes in mRNA expression and thereby cellular functions^1,2^. Inside the crowded environment of the cell, distinct extracellular signals frequently use common effector proteins (kinases) serving as intersections (cross-talk points) between pathways. Hence, signaling pathways relaying information through the cell must be approached as complex interconnected networks, rather than isolated silos. A critical implication of this view is that the route the initial signal takes through these networks governs which nuclear factors are impacted and, therefore, the transcriptional response. As such, the ‘wiring’ of this network (e.g. which substrates are available to the activated kinases and their connections) determines the possible paths available for the signal to regulate cellular processes.

Most of our current understanding of signaling pathways and their connectivity is based on cell population averages and qualitative methods. This established Epidermal Growth Factor (EGF) Receptor signaling as an interconnected network of kinases, containing multiple distinct pathways with different functions. Two of the major signaling pathways in the EGF signaling network are the Akt-mTOR and Ras/ERK (MAPK) signaling cascades^1,2^. Both these signaling routes lead to phosphorylation of ribosomal protein S6 (RPS6), as well as downstream DNA-binding transcription factors^1,3^. For instance, the p90RSK kinase activates RPS6 downstream of MAPK signaling through Erk1/2, whereas p70S6K phosphorylates RPS6 upon activation by Akt-mTOR ^2^. Importantly, many potential points of cross-talk and feedback have been described between EGF– and other signaling pathways. This is exemplified by the inactivating phosphorylation of Wnt-pathway component GSK3β (on Serine 9) by Akt/PKB^1,2^, and activation of JAK-STAT signalling component STAT3 via RAS/ERK signaling^4^. Recent years have seen advances in studying signaling dynamics using single-cell approaches. For instance, mass cytometry analysis of overexpressed GFP-tagged kinases revealed an interdependence of kinase effector abundance and the dynamics of signaling progression through the EGFR pathway^5,6^. Additionally, optogenetic activation of Son-of-Sevenless (SOS) allowed quantitative screening of drug treatment-effects on the Erk/MAPK-pathway, demonstrating that targeted drugs can corrupt its dynamic signal-transmission properties in a potentially pathological manner^7^.

Cancer-associated mutations are often found in components of key developmental signaling pathways, which has led to a great interest and investment in developing small molecule drugs against specific kinases to block these cancer driving pathways^8,9^. Although promising and powerful, the emergence of drug resistance is a major problem in current clinical practice^10–13^. Identified drug resistance mechanisms include the selection/appearance of cells with desensitizing genetic mutations, as well as the existence of seemingly mutation-independent stochastic drug-tolerant cell-states^14–17^. A common denominator of drug resistance mechanisms is that the way the extracellular signals are perceived and processed is changed to benefit the resistant cell, indicating that variation in the signaling network can greatly impact the cellular outcome. The observation that even within seemingly homogenous cell populations not all cells respond in the same way, suggests that intrinsic (or ‘cell-state’) differences between cells influence their response ^18,19^. Cell-states include, for instance, the combination of prior and current signaling events, cell-cycle status, differentiation level and position with respect to other cells. How the internal state of a cell affects the flow of information inside the cell and the subsequent transcriptional response is still largely unknown. Considering that the response of a given cell to an environmental signal is inherently cell-autonomous (i.e. it is determined by intracellular molecular mechanisms), the processes involved must be studied at the level of individual cells.

Here, we describe the use of cultured primary human epidermal keratinocytes to study drug-responses at the resolution of individual cells using single-cell ImmunoDetection-by-sequencing (scID-seq) to measure ∼70 (phospho-)proteins per cell in different conditions. Using the core Epidermal Growth Factor Receptor signaling pathway as a paradigm, we found that blocking down-stream pathway effectors p90RSK and p70S6K resulted in cell-state dependent signaling responses. Using a newly developed single-cell Comparative Network Reconstruction (scCNR) model, we identified the differences in signaling network wiring associated with these cell-states.

## Results

### Single-cell (phospho-)protein profiling of the response to p90RSK and p70S6K inhibition

We used primary human foreskin keratinocytes as a karyotypically normal and genetically stable cell system for our experiments to avoid confounding effects of heterogeneity with respect to cancer-associated mutations in signaling components. To capture how signals are processed, and how cells respond to perturbations, we subjected keratinocytes to short term inhibition of key effectors of the MAPK and AKT-mTOR pathways downstream of the EGFR (Figure 1a). We reasoned that focusing on early effects would allow us to associate pre-existing cell-to-cell heterogeneity with differential cellular responses. Keratinocytes were grown in medium without exogenous EGF for 24 hours prior to the experiment, in which we treated the cells for 2 hours with an inhibitor of p90RSK (BI-D1870, 10 μM) or p70S6K (PF4708671, 10 μM), or a vehicle control (DMSO), followed by a 30-minute stimulation with 10 ng/ml EGF while maintaining the presence of the inhibitors. After harvesting and fixation, we applied scID-seq to measure the abundance of ∼70 (phospho-)proteins in each cell using specific DNA-barcoded antibodies. Our panel contains characterized and validated antibodies detecting phosphorylated and non-phosphorylated components from the Akt-mTOR signaling and MAPK signaling, as well as other key developmental and cancer related pathways (Wnt, BMP, TNFα, TGFβ, Notch, Jak/STAT and Integrin-mediated adhesion) and cellular processes including the cell cycle and differentiation ^20–22^. We used cell-hashing^23^ as a sample barcoding approach to minimize batch-effects and variation in staining efficiencies between conditions (Figure 1b and TableS1). After pooling the samples and staining with the full antibody panel, individual cells were distributed into 96-well plates using FACS, effectively randomizing the treatment groups over the different independent sample preparations that are performed per plate. After sequencing, quality control, pre-processing and filtering (see Methods section for more information), we obtained a dataset consisting of 69 (phospho-)protein measurements from 546 cells (Table S1).

**Figure 1.**
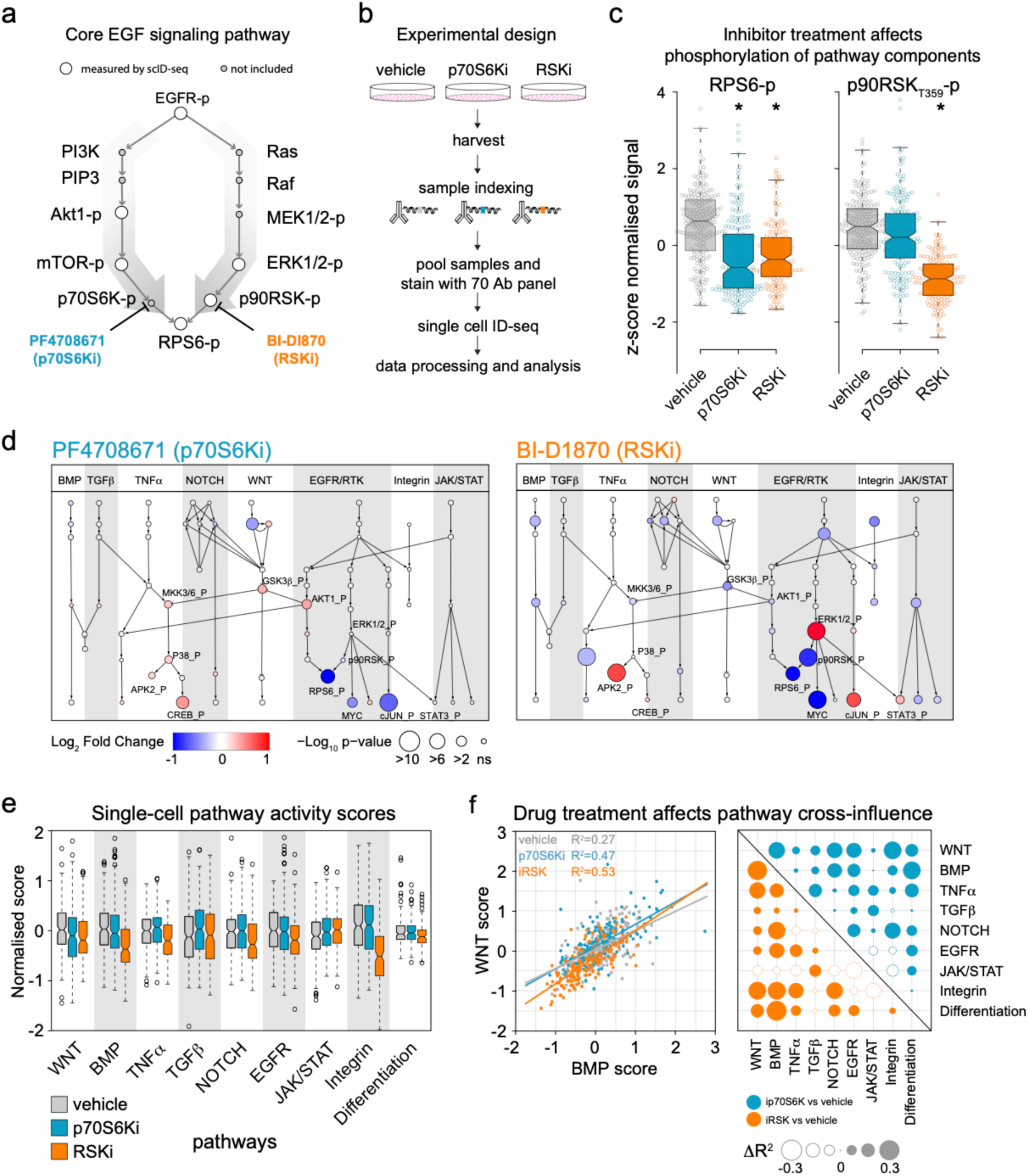
Single cell ID-seq profiling of signaling protein activity shows widespread and heterogeneous response to targeted inhibition. **a.** Schematic overview of the AKT and MAPK signaling pathways. **b.** Schematic overview of the scID-seq workflow. **c.** Boxplots of phospho-RPS6 (left) and phospho-p90RSK abundance upon drug treatment. Abundances are TTM-normalized and subsequently z-score transformed to facilitate comparison. **d.** Differential abundance of signaling phospho-proteins upon p70S6Ki (left) and RSKi (right) shows that treatment effects propagate throughout the signaling network. Colors indicate log_2_-fold change, node sizes indicate –log_10_ p-values. Edges between proteins are drawn based on literature-established interactions. **e.** Single cell pathway activity scores upon drug treatment. Pathway activity scores are obtained by combining multiple features that belong to a pathway, taking into account whether they have an activating or inhibiting effect on the pathway. **f.** Correlation between the pathway activity scores, calculated over cells within a treatment.

### Drug treatment leads to alternative signaling flow and transcriptional responses

Consistent with their known function of phosphorylating RPS6, inhibition of either p90RSK (BI-D1870) or p70S6K (PF4708671) resulted in pronounced reduction of phospho-RPS6 levels, whereas p90RSK-T359 phosphorylation specifically decreased upon p90RSK inhibition, providing confidence in our experimental set-up and scID-seq measurements (Figure 1c). Inspection of further core EGFR pathway components revealed that ERK1/2 and AKT1 phosphorylation signals increased upon p90RSK and p70S6K inhibition, respectively (Figure S1a). This is consistent with their involvement in a regulatory negative feedback loop, a well-known design principle to control signaling pathway activity ^24,25^, and raises the question whether these perturbation effects propagate through the wider signaling network. We investigated whether these accumulated phosphorylation levels were associated with activation of other pathways through so-called signaling cross-talk. Differential abundance analysis revealed significantly changed (phospho-)protein levels for multiple components of the JNK, p38 and Wnt pathways in cells treated with the p70S6K inhibitor (Kolmogorov-Smirnov test, Figures S1b, S2 and Table S2). Notably, GSK3β Serine-9 phosphorylation, a known pathway cross-talk event catalyzed by Akt ^1,2,26^, was increased. In turn, inactivation of GSK3β was associated with enhanced phosphorylation of MKK3/6 (Figure 1d, S1a and S2, see Figure S3 for all node identities), consistent with previous findings ^27,28^. Furthermore, phosphorylation levels of a known downstream effector of MKK3/6, MAPK-p38, and its subsequent targets MAPK-APK2 (henceforth APK2) and the CREB transcription factor ^29,30^ were also significantly increased (Figure 1d, S1a and S2). Similarly, p90RSK inhibition stimulated phosphorylation of STAT3 and decreased phosphorylation of Jak1 and STAT1 (Figure 1d, S1a and S2, Table S2). Phosphorylation of STAT3 by MAPK signaling was previously described as a cross-talk mechanism ^31,32^. Additionally, we found increased phosphorylation of APK2, independent of its canonical upstream kinases MKK3/6 and MAPK-p38 in p90RSK inhibitor treated cells (Figure 1d, S1a and S2). Rather, this APK2 phosphorylation may be a consequence of accumulated ERK activity through direct phosphorylation of the Thr334 site on APK2, as previously described ^33^. Together, these data indicate that p70S6K and p90RSK inhibition resulted in accumulated phosphorylation of upstream regulators (Akt and ERK, respectively), an effect that propagates into alternative pathways in the biochemical signaling network through cross-talk points.

So far, we only considered the mean abundance of each (phospho-)protein per treatment, but there is clear within treatment variation between cells (Figure 1c). To investigate interactions between pathways more coherently, we derived aggregated activity-scores for each pathway in each cell (see Methods for details). We confirmed that p70S6K and p90RSK inhibitors indeed differentially affect the signaling pathways covered by our antibody panel (Figure 1e). Although such aggregate scores for pathways cannot fully account for all regulatory intricacies and complexities, it does allow us to systematically assess interactions between pathways while extending it to situations where the points of cross-talk within the signaling network are unknown, and/or not directly represented in our measurements. To investigate pathway cross-influence, we quantified the proportion of the variation *within* a treatment condition in a given pathway (i.e. the response variable) that can be explained by the variation in another pathway (i.e. the explanatory variable) by calculating the coefficient of determination (R^2^) between pathway scores across individual cells for each of the treatment conditions. For example, the coefficient of determination between the BMP and Wnt pathways in vehicle treated cells was R^2^=0.27 (i.e. 27% of variation was explained), whereas this increased to R^2^=0.53 upon p90RSK inhibition, almost doubling the proportion of variation explained (Figure 1f, left panel). Performing this analysis for all pathway combinations and expressing the difference in the coefficient of determination between vehicle and treatment (ΔR^2^), revealed evidence of strong and pervasive perturbation-dependent effects on pathway cross-influence (Figure 1f, right panel). These analyses indicate that treatment with targeted inhibitors can lead to wide-spread propagation of their effects to other pathways within the biochemical signaling network that could shape the cellular response.

A key mechanism by which extracellular signals and their intracellular transduction pathways influence these responses, and consequently the biology of the cell, involves regulating (sets of) DNA-binding transcription factors. Therefore, a potential consequence of the observed differential cross-talk is that a different complement of transcription factors is regulated, leading to altered transcriptional output. Consistent with this, p90RSK inhibition resulted in differential (phosphorylation) levels of the transcription regulators Myc and c-Jun (Figure 1d). To investigate the consequences on mRNA expression, we treated keratinocyte cultures with the p90RSK inhibitor BI-D1870 (10 μM) for 2 hours followed by single-cell RNA-sequencing. After processing and normalization, clustering and t-SNE representation, we found that vehicle and p90RSK inhibitor treated cells separate into distinct groups based on their RNA profiles (Figure S1c). Moreover, c-JUN mRNA levels were increased, consistent with the (phospho-)protein data and the fact that c-JUN engages in an autoregulatory positive feedback loop ^34^. The early growth response gene 1 (EGR1) transcript, a canonical c-JUN target, is simultaneously upregulated, indicating that c-JUN transcription regulation is affected by p90RSK inhibition. Moreover, differential expression testing between the two treatments and motif analysis identified enrichment of AP1 (c-JUN) and Myc binding sites in promoters of deregulated transcripts (Figure S1d, left panel). These observations were confirmed with an independent p90RSK inhibitor (LJH-685, 10 μM, 2 hours, Figure S1d, right panel). These findings indicate that even though treatment with a drug blocks the intended target, it can result in activation of alternative pathways through the biochemical signaling network, and unintended transcriptional consequences.

### Robust identification distinct cell-state clusters from scID-seq data

A key implication of the findings described above is that the route a signal takes through the network is influenced by whether the signal can be efficiently transmitted to the next node (i.e. network wiring), which is in essence the process that was disrupted by the inhibitor treatment. We hypothesized that this wiring may vary depending on which *cell-state* the cells reside in. Such cell-states can include the position in the cell cycle, the degree of differentiation, ongoing or prior signaling events, and other processes. If this hypothesis is correct, this should be reflected as structured cell-to-cell variation in (phospho-)protein levels in our dataset. Indeed, we observed clear effects on the average phosphorylation level that were accompanied by large within-treatment variations for many of the included (phospho-)proteins (Figure 1c, S1a, S2), hinting at the existence of multiple cell-states in our data.

Identifying cell-states can be challenging due to confounding and dominating effects associated with the applied treatment. Inspired by previous work from Kramer and Pelkmans ^18^, we defined 23 (phospho-)proteins that did not show strong treatment responses (Kruskal-Wallis test, comparing vehicle with p70S6Ki and p90RSKi, –log_10_ p-value <5, c.f. Table S2) as ‘cell-state markers’, and designating all other (treatment responsive) (phospho-)proteins ‘signaling-state markers’. Subsequently, we developed a robust cluster assignment pipeline, based on the Nearest Neighbor (NN)-dependent Leiden clustering algorithm and identified 9 distinct clusters, which were superimposed on a UMAP representation of our scID-seq dataset (Figure 2a) ^35^. It is important to note that the Leiden algorithm is non-deterministic in nature as it contains a random NN-network initialization step, leading to unstable cluster calling in repeated iterations (Fig S4a). Consequently, cells on the ‘borders’ between clusters can be mis-assigned. To solve this, we adapted the workflow into a consensus/ ‘majority vote’ approach, using 1000 random initializations to identify the most frequently occurring (i.e. stable) number of clusters (Figure S4a). This is then repeated 5 times and the level of co-assignment of cells in the same cluster is determined (Figure S4b,c). Using this approach >98% of the all cells were unambiguously assigned (i.e. consistently assigned into the same stable cluster in all 5 iterations, Figure S4d). Moreover, the treatment groups were relatively well distributed over the different clusters, indicating that cluster assignments were not strongly influenced by treatment effects (Figure S4e). Some of these cell-states are associated with specific cell-cycle stages, as evidenced by their levels of Cyclin B1 and Histone H3S10 phosphorylation (Figures 2b and 5e), whereas other clusters express keratinocyte basal/progenitor cell marker TP63 (Figure 2b), allowing a degree of biological interpretation to the identified cell-state clusters. Therefore, the 9 stable clusters we identified reflect underlying cell-states prior to treatment.

**Figure 2.**
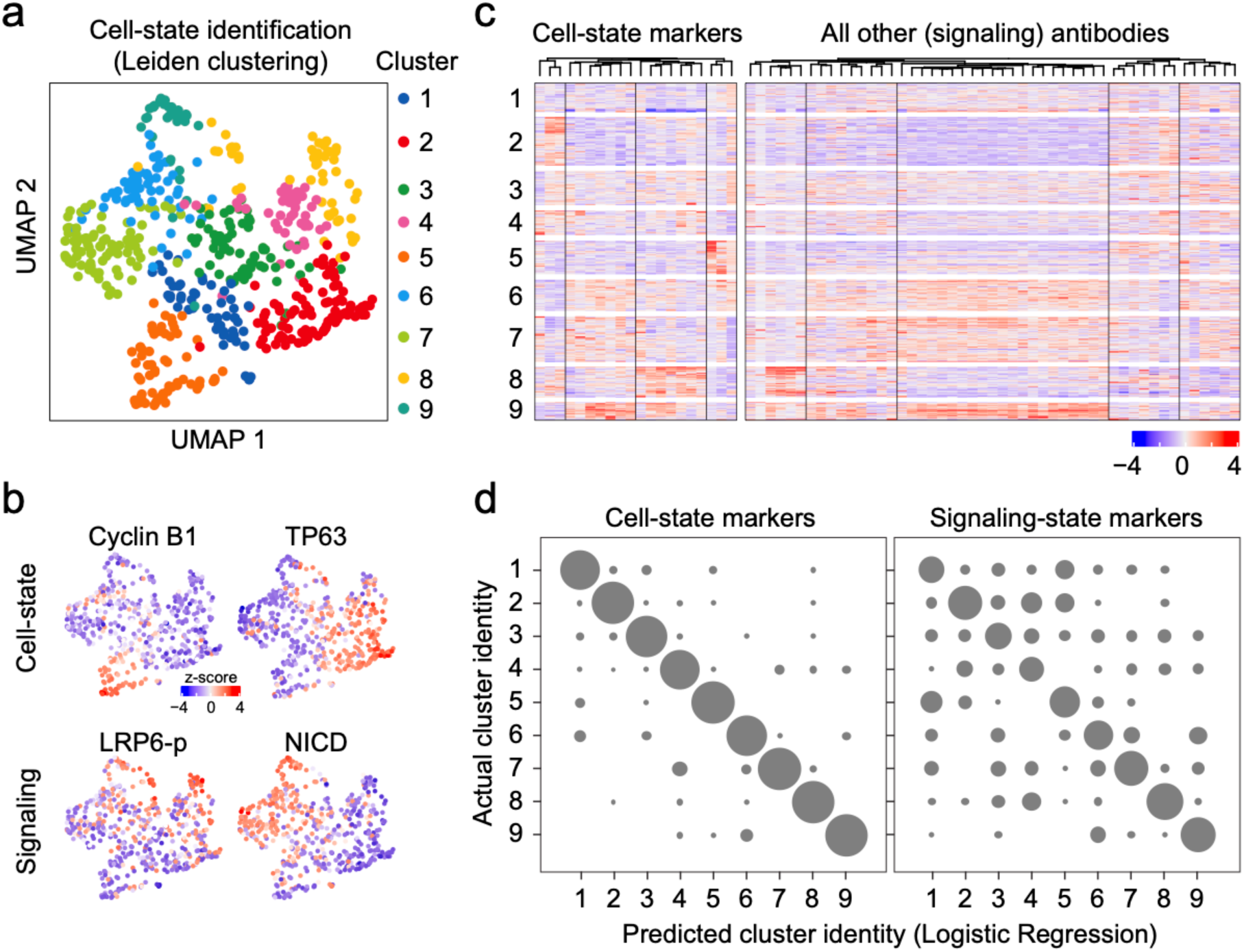
Robust identification of nine distinct cell-states. **a.** UMAP projection of the cells, color coded according to their cell-state assignment. Only the cell-state markers (i.e. (phospho-)proteins that did not show a differential abundance between treatments) were used in calculating the UMAP coordinates. **b**. UMAP projection of the cells, color coded by the abundance of selected cell-state (top row) or signaling-state (bottom row) markers **c.** Heatmap of (phospho-)protein abundances across cells. Rows indicate cells, grouped by cell-state, columns indicate (phospho-)proteins, grouped by cell-state or signaling-state markers. The signaling-state markers, which were not used in the cell-state identification, nonetheless show strong cell-state dependent expression patterns. **d.** Multiclass confusion matrix showing the performance of a logistic-regression classifier in predicting the cell-state cluster identity of cells, based on cell-state markers (left) or signaling-state markers that were not used in the cell-state assignment (right). Performance was assessed in a 5-fold nested cross-validation loop.

To explore potential associations between these cell-states and signaling activity we visualized the dataset in a heatmap grouped by cell-state, re-affirming their cluster-associated expression patterns (Figure 2c). Interestingly, the signaling-state markers also exhibited strong cluster dependent patterns (e.g. phospho-LRP6 and Notch Intracellular Domain (NICD)), even though these (phospho-)proteins were not used for cell-state identification (Figures 2b,c). This indicates that the identified cell-states capture biological distinctions that are also associated with specific signaling pathway activities. To quantify the extent to which the signaling markers contain information about the cell-state in a more objective way, we trained Machine Learning classifiers (Multinomial Logistic Regression (MLR), Random Forest (RF) and Support Vector Machine (SVM)) to “predict” the cell-state from the signaling-state markers only. As a control we trained these classifiers on the cell-state markers that were used in the original cell-state assignment. In a nested 5-fold cross-validation loop, MLR reached ∼88% classification accuracy (Figure 2d). RF and SVM classifiers reached average accuracies of ∼81% and ∼87%, respectively (Figures S5a-c). When trained on the signaling-state markers in the same nested cross-validation setting, MLR consistently obtained 51% accuracy of predicting the true cell-state labels (Figures 2d and S5c). In contrast, an MLR classifier trained on data with randomized labels only reached ∼12% accuracy, as expected based on chance (Figure S5c). Similar results were obtained using RF and SVM classifiers (Figure S5b-c). Together, these analyses show that the identified cell-states contain additional intrinsic underlying differences in signaling activities that potentially influence eventual treatment effects of cells in these states.

### Cell-state dependent drug responses are pervasive and specific

Our experimental design and the analyses thus far enable us to investigate cell-state dependence of treatment responses. For instance, if the effect of inhibiting p70S6K or p90RSK is independent of cell-state, we would expect treatment-effects to be the same, or at least highly similar, across the identified clusters. In contrast, we observed a high degree of variation of the effects sorted by either inhibitor among the cell-state clusters for individual (phospho-)proteins, and at the level of pathway activity scores (Figures 3a and S6a,b). To statistically test the extent to which the signaling markers show cell-state dependent responses to the drug treatments, we fitted a linear regression model that aims to explain (phospho-)protein abundance using drug-treatment, cell-state, and an interaction term between these, for each measured (phospho-)protein. In such models, the interaction term quantifies the extent to which the abundance of a (phospho-)protein in a cell differs from what would be expected based on the drug perturbation and the cell-state separately. We assessed the significance of the interaction term both by using an ANOVA and by considering the coefficients from the linear model directly (Figure 3b and Table S3). The latter approach has the benefit that each interaction between each cell-state-drug treatments is assessed, but has the limitation that it requires a reference for the linear regression analysis. Although the vehicle control could be used as a logical reference for the treatment, there was no obvious reference with regards to cell-state. We opted to use cell-state 1 as the reference, and confirmed that using any of the other cell-states as reference yielded consistent results (Figure S7a). This analysis revealed 100 significant cell-state dependent treatment responses involving 50 out of the 69 analyzed (phospho-)proteins, across all 8 cell-states (other than the reference cell-state 1) and both inhibitors (p-value<0.05, Figure 3b, Figure S7b and Table S3). Examples include the specific upregulation of cyclinB1 and downregulation of histone H3S10 phosphorylation levels in cluster 5. Additionally, Cyclin E displays a strong positive interaction between p70S6K inhibition and cell-state 9 (Figure 3b,c). In contrast, ERK1/2 phosphorylation is an example of a phospho-protein that shows cell-state independent, inhibitor treatment specific response (Figure 3b,c). To examine whether cell-state dependent responses propagate through the signaling network, we returned to the analyses of pathway cross-influence based on the activity scores and systematically calculated the change in coefficient of determination among pathways between vehicle and treated cells within each cell-state cluster. This revealed major differences in how pathways influence each other across the different clusters (Figure S6c). Overall, our analyses identified cell-state specific drug-responses that may result from underlying differences in signaling network wiring.

**Figure 3.**
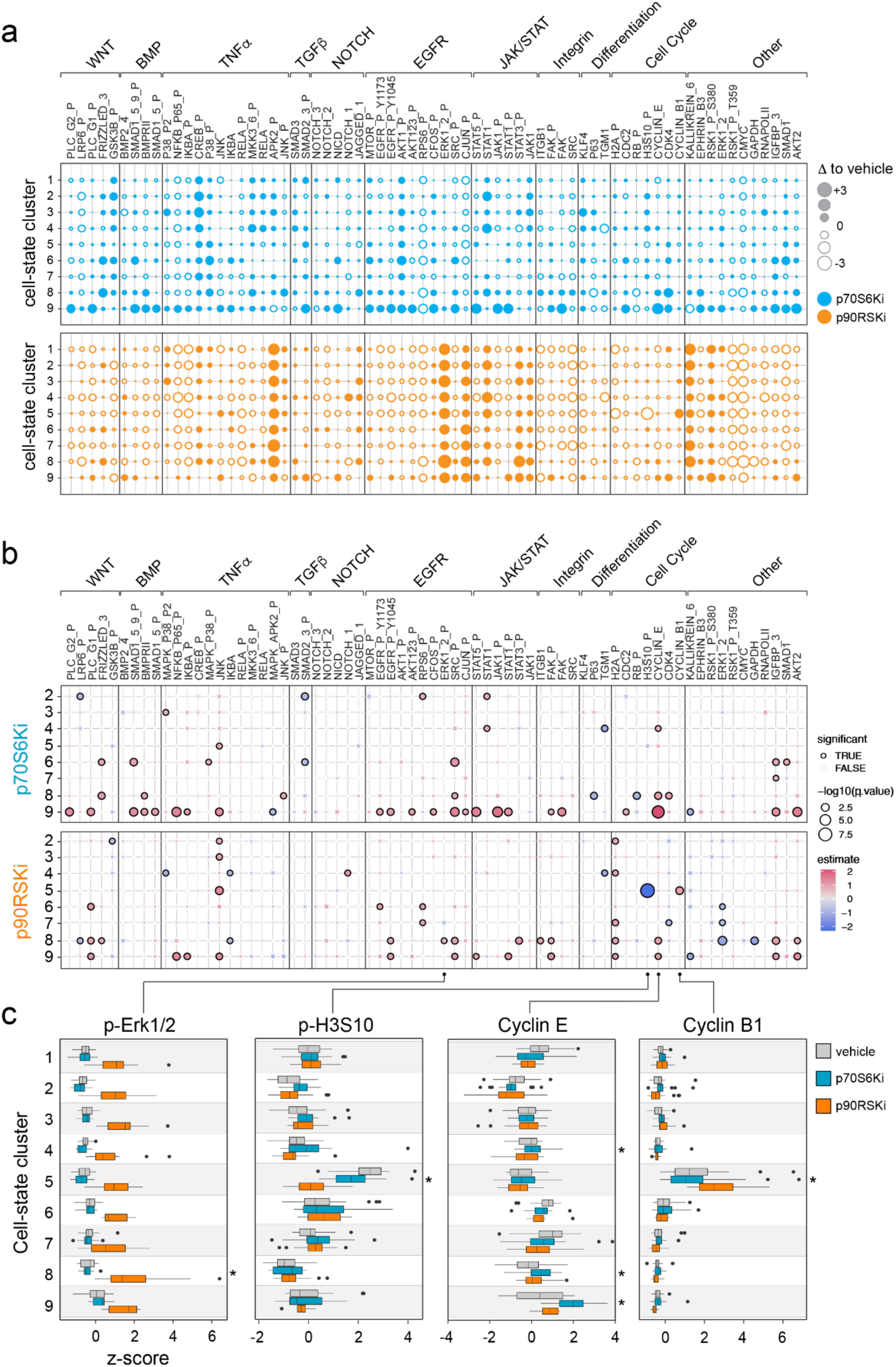
Cell-state specific responses are pervasive and drug dependent. **a.** Differential (phospho-)protein abundance in cells treated with vehicle versus p70S6Ki (blue) or p90RSKi (orange). Open and closed circles indicate down– and upregulated levels, respectively. **b.** Overview of cell-state specific drug response for all cell-state (phospho-)protein pairs, as assessed by a linear model with an interaction term, where cell-state 1 was used as a reference. Colors indicate the effect size of the interaction term coefficient, point size indicates the –log_10_ p-value. Significant interactions (p-value < 0.05) are indicated by a black outline. **c.** Boxplots of the abundance of Cyclin B1, Cyclin E, phospho-ERK1/2 and phopsho-H3S10, split out per treatment and cell-state cluster, to exemplify cell-state dependent drug-responses. Cell-states with significantly different drug-responses are indicated with an asterisk.

### Single-cell Comparative Network Reconstruction reveals cell-state specific signaling flow and network wiring

To quantitatively investigate differences in signaling network wiring between cell-states, we recently developed single-cell Comparative Network Reconstruction (scCNR). scCNR is designed to exploit the within cell-state variation in signaling activities of the network nodes to quantify their interaction strengths and is described in more detail elsewhere ^36^. In short, the interaction strength r_ij_ is defined as the percent change in activity of node i upon a 1% change in activity of node j, if all other nodes were to be kept constant. scCNR fits a model with the same topology (i.e., which interactions are present and absent) for each cell-state, but allows for differences in interaction strengths between states. Importantly, our method can take prior information about network topology as an input, but can also suggest additional interactions if these significantly improve the model fit. By penalizing the number of interactions in the network, a balance between model complexity and data fit is ensured. Similarly, by penalizing the number of interactions that differ in strength between the cell-states, the most relevant differences in the signaling network between the cell-states are obtained. First, we established a topology for our network based on literature-derived canonical interactions (see also Figure S3) and interactions obtained from the public phosphosite plus database ^37^. Subsequently, we varied the penalty on model complexity to examine the extent to which the model fit could be improved by including extra edges.

Adding 34 edges resulted in a good balance between model fit and complexity (Figure S8a,b). To validate the output topology from our model, we compared it to 1000 models with random topologies (but with the same total number of edges) and quantified model performance as the root mean square of residuals (RMSR). The RMSR of our network model is significantly smaller and falls well outside the distribution of RMSRs from the 1000 random models (p-value < 0.001, Figure S8c).

Having established the performance of our network reconstruction model, we set out to identify network interactions whose strength differs between cell-states. Using a penalty on the difference between cell-states that gives a sensible balance between number of differences and model fit (Figure S8d), our scCNR algorithm highlighted 31 edges that account for most of the cell-state specific signaling flow (Figure 4a). Combined, these 31 cell-state specific interaction strengths attain 79% of the total reduction in RMSR that can be gained from cell-state specific edges (Figure S8d). We assessed the robustness of the identification of cell-state specific edge strength using a bootstrapping experiment, which showed that the majority of cell-state specific edges are identified in the majority of all bootstraps (Figure S8e). To identify for which edge-cell-state combinations these effects are most important, and to assess the statistical significance of these differences, we performed permutations analyses in which we randomized cell-state assignments while preserving the number of cells from each treatment in each cell-state. We then optimized the edge-strengths while retaining the network topology, and fixing which edges differ between cell-states. We repeated this procedure 1000 times to obtain a null-distribution and an empirical p-value of the cell-state dependence of each edge in the network. This identified 22 significant cell-state edge pairs that were different between cell states, which were fairly evenly distributed over the different cell-states (FDR <0.05, Table S4). The notion that cell-state, drug response, and network effects are interwoven, is exemplified by the response of Cyclin B1 to p90RSKi. As discussed above, the inhibition of p90RSK resulted in accumulated phospho-Erk1/2 levels, presumably through a negative feedback loop, which in turn leads to APK2 activation (Figure 1d, S1a and S2). Our network reconstruction results expand on this and identified a cell-state dependent interaction where APK2 activation explains Cyclin B1 accumulation specifically in cell-state 5, but not in any of the other cell-states (Figures 4c,d and S8f). Indeed, high levels of cell-cycle markers phospho-H3S10 and CyclinB1 in vehicle-treated cells confirm that cells in state 5 are undergoing G2/M transition (Figures 3c and S5d). Consistent with this, the mitosis specific marker phospho-H3S10 ^38^, selectively fails to accumulate in cell-state 5 when p90RSK is inhibited (Figure 3c). Although p90RSK2 was historically described to phosphorylate H3S10 downstream of EGF signaling independently of mitosis ^39^, our findings show that p90RSK-mediated APK2 activity is required for human keratinocytes to efficiently complete the G2/M transition.

**Figure 4.**
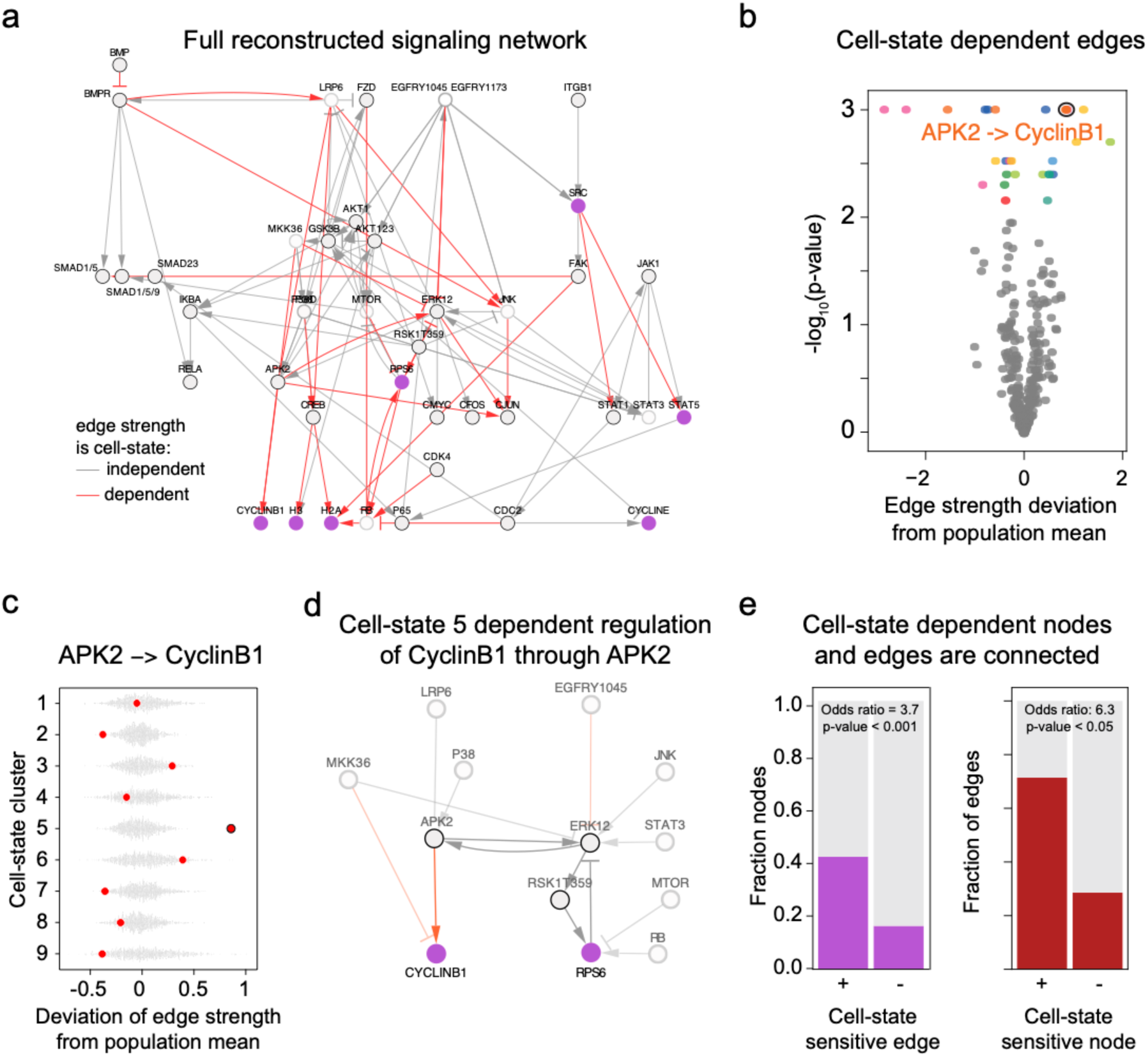
Cell-state specific signal transduction and drug response are tightly connected. **a**. Full reconstructed signaling network. Red edges indicate Interactions that quantitatively differ between cell-states. Purple nodes are (phopsho-)proteins that show cell-state specific drug response. **b**. Volcano plot showing the –log_10_p-value and interaction strength deviation from the population mean for all edges that were reconstructed to be cell-state specific (red edges in panel a) and all cell-states. The null distribution was obtained by randomly permuting the cell-state labels and reoptimizing the interaction strengths. Interaction-cell-state pairs that are significantly different in a particular cell-state from the population mean are highlighted color, corresponding to the same color code as in Figure 2a. **c.** Example of an edge (from APK2 to Cyclin B1) whose strength deviates significantly from the population mean, specifically in cell-state 5. Red points indicate the true reconstructed edge strengths, gray dots represent strengths from reconstructions where the cell-state labels were randomly permuted. In most cell-states, the interaction from APK2 to Cyclin B1 is nearly absent, but in cell-state 5 it is a strong, positive interaction. **d.** Highlighted part of the full network that explains the strong cell-state dependence of Cyclin B1 expression. The reduction in RPS6 upon RSKi causes feedback activation of ERK1/2, which in turn activates APK2. Due to the cell-state specific interaction between APK2 and Cyclin B1, only in cell-state 5 does this activation of APK2 translate into an accumulation of Cyclin B1. **e.** Visual representation of the contingency tables describing the connection between cell-state dependent drug response and interaction strengths. Left panel: The bars indicate the fractions of the nodes that show cell-state dependent drug response, separated by whether or not they have an income edge that has a cell-state dependent strength. Right panel: Fraction of edges that have a cell-state dependent strength, separated by whether or not they connect (upstream or downstream) to a node showing cell-state dependent drug response.

Importantly, these observations indicate that cell-state specific differences in node response and edge strength (i.e. network wiring) may be closely related. To test this hypothesis, we investigated if edges that differ between cell-states are more likely to be connected to nodes that show cell-state dependent drug response throughout the whole network. Indeed, 42% of edges that differ between the cell-states are connected to nodes with a significant interaction term, whereas only 16% of edges that do not differ between cell-states are, representing significant enrichment (Figure 4e left panel, Fisher exact test: odds ratio = 3.7, p<0.001).

Conversely, 71% of nodes with a significant interaction term receive input from an edge that has a cell-state specific strength, in contrast to only 29% of nodes without a significant interaction term (Figure 4e right panel, Fisher exact test: Odds ratio = 6.3, p<0.05). These results also hold in more fine-grained cell-state specific analyses (Figure S8g). Taken together, our scCNR approach revealed that quantitative differences in signaling network wiring are associated with cell-state dependent drug responses.

## Discussion

The question of what makes some cells respond to a given signal differently from other cells is central to understanding processes in normal and disease biology, ranging from symmetry breaking and patterning in early development, to the emergence of therapy-resistance in cancer. We approached this question by characterizing intracellular signaling network activities and wiring upon small molecule inhibitor treatment at the level of individual cells using single-cell ImmunoDetection by sequencing (scID-seq). This work revealed several underappreciated aspects of cellular responses to such treatments. First, inhibiting a downstream effector kinase may result in accumulated activity of upstream pathway components, presumably through disruption of feedback loops. Second, this signal can propagate into alternative routes through the intracellular biochemical signaling network, leading to unintended regulation of downstream DNA-binding transcription factors and their subsequent transcriptional programs. Third, we used the multidimensionality of our data to define 9 distinct cell-states present in our cell population, and found that the response to the inhibitor treatment strongly depended on which of these states a cell resided in. Fourth, using a newly developed single-cell Comparative Network Reconstruction method ^36^, we were able to identify the quantitative differences in signaling network wiring that are associated with these cell-state dependent drug-responses.

Our work is consistent with previous findings on cell-type differences in oncogenic signaling in colorectal cancer ^40,41^ and with the multimodal perception concept put forward by Kramer and Pelkmans ^18^. Together this indicates that redirected and cell-state dependent signal processing may be inherent to the mechanism of action of targeted therapies, and that these effects need to be accounted for when designing therapeutic strategies. One way to achieve this is to devise strategies to coax cells into cell-states in which they will exhibit the desired response to treatment ^42^ that can take the form of neoadjuvant, or combination therapy regimes. However, this will require detailed understanding of cell-state differences in network wiring, as well as computational models to identify suitable perturbations to achieve this ^43^.

The approaches and results we presented here can provide a framework for such future directions. Additionally, the fact that our work was performed in primary human cells in the absence of transformation by cancer-associated mutations indicates that cell-state dependent cellular responses are not restricted to cancer-contexts and may also shape normal biology.

## Methods

### Cell culture

Primary pooled human epidermal stem cells derived from foreskin were obtained from Lonza. Cells were cultured and expanded as previously reported ^44^. Briefly, cells were cultured on a feeder layer of J2-3T3 cells in FAD medium (Ham’s F12 medium/Dulbecco’s modified Eagle medium (DMEM) (1:3) supplemented with 10% batch tested fetal calf serum (FCS) and a cocktail of 0.5 μg/ml of hydrocortisone, 5 μg/ml of insulin, 0.1 nM cholera enterotoxin, and 10 ng/ml of epidermal growth factor) supplemented with Rock inhibitor (Y-27632, 10 μM). J2-3T3 cells were cultured in DMEM containing 10% bovine serum and inactivated with MitomycinC (SCBT) upon seeding the epidermal stem cells. For experiments epidermal stem cells were transferred to Keratinocyte Serum Free Medium (KSFM) supplemented with 0.2 ng/ml Epidermal Growth Factor (EGF) and 30 μg/ml bovine pituitary extract from Life Technology until 70% confluency. Cells were treated with p90RSK inhibitor BI-D1870 (10 μM, Sellekchem) or P70S6K inhibitor PF4708671 (10 μM, Sellekchem). All media were supplemented with 1% penicillin/streptomycin antibiotics. We included a non-stimulated control and found that endogenous EGFR ligands produced by the cells themselves circumvented the need for exogenous pathway activation (data not shown). Although this control group was omitted from further analyses, these measurements are included in the full dataset.

### Antibody conjugation with dsDNA barcodes

The antibody panel, including extensive validation, dsDNA functionalization and conjugations were performed as described^20,21,45^. In short, antibodies were functionalized with NHS-s-s-PEG4-tetrazine (Jena Bioscience) in a ratio of 1:10 in 50 mM borate buffered Saline pH 8.4 (150 mM NaCl). Then, N3-dsDNA was produced and functionalized with DBCO-PEG12-TCO (Jena Bioscience) in a ratio of 1:25. Finally, purified functionalized antibodies were conjugated to purified functionalized DNA by 4-hour incubation at room temperature in borate buffered saline pH 8.4 in a ratio of 4:1 respectively. The reaction was quenched with an excess of 3,6-diphenyl tetrazine. The conjugation efficiency and quality were confirmed on an agarose gel and equal amounts of each conjugate were pooled for scID-seq experiments.

### Immuno-staining and single cell sorting

Staining and sorting was performed as described previously. Briefly, Cells (> 3 x 10^6^) were harvested with trypsin and cross-linked in suspension by incubating for 10 minutes with 4% paraformaldehyde (PFA) in PBS following a quenching step of 5 minutes with 125 mM Glycine in PBS. Removal of PFA and Glycine occurred through washing twice with a wash buffer (0.1x Pierce^TM^ Protein-Free Blocking Buffer from Thermo in PBS). Then, cells were blocked in 500μl blocking buffer (0.5x Pierce^TM^ Protein-Free Blocking Buffer, 200 μg/ml boiled salmon sperm DNA, 0.1% Triton-X 100, in PBS) at room temperature for 30-60 min. Pre-stainings were performed at room temperature for 1-2 hours and staining with the conjugate mix occurred overnight at 4°C in 500 μl blocking buffer. After each staining, cells were washed 3x in 5 ml wash buffer. Cells were sorted in single wells with the BD FACSAria SORP flow cytometer (BD biosciences) in 96 well PCR plates containing 1 μl release buffer (10 mM DTT in 15mM Tris, pH 8.8) and 7 µl Vapor-lock (Qiagen).

### Barcoding and library preparation for next generation sequencing

#### scID-seq

Library preparation was performed as described previously^21^. Briefly, 3 PCR steps were performed to amplify the antibody barcodes and to add barcodes specific for the well and the plate of each cell. The barcoding occurred with the same sequences used in ID-seq ^20^. For the first PCR step 15 cycles were run after adding to each well a 4 µl reaction mix containing the Herculase II Fusion DNA Polymerase (Agilent), dNTPs, 5x Herculase buffer and 0.1μM amplification primers. Directly after the first PCR step, 5 extra cycles were run after adding 1 µl mix containing Herculase buffer 0.2 μM forward amplification primer and 0.2 μM reverse well barcoding primer. Then all material was pooled per plate, Vapor-lock was removed and a clean-up was performed with the QIAquick PCR Purification Kit, followed by an exonuclease 1 treatment to degrade remaining primers and another purification. Another 5 cycles were run in PCR 3 with a 20 µl reaction containing the pooled and purified sample and 0.1 μM plate-barcoding primers. After repeating the clean-up procedure, the libraries were checked on agarose gel and with the Bioanalyzer (Agilent) to confirm the size of the DNA fragments (expected size around 185 bp) and sequenced on an Illumina NextSeq500 machine.

#### scRNA-seq

Library preparation was performed as described previously^46^. Briefly, cultured cells were dissociated, followed by fixation with 2.5 mM DSP and 2.5 mM SPDP in sodium phosphate buffered saline (pH 8.4) for 45 minutes at RT n order prevent transcriptional changes during the single-cell sorting, The fixation was quenched by adding 100 mM Tris-HCl pH 7.5 and 150 mM NaCl for 10 minutes. Individual cells were sorted (BD FACS Aria) into 384-well plates containing CEL-seq2 primers. The cDNA library preparation was adapted from the previously published protocol^46^ and ERCC spike-in (1:100,000 dilution, Thermo Fisher Scientific) was added to the reverse crosslinking mix. The cDNAs were selected using a 0.6x Ampure XP bead ratio (Beckman Coulter) purification and subsequently subjected to second strand synthesis and subjected to *in vitro* transcription based amplification using Megascript T7 transcription kit (Thermo Fisher Scientific). The amplified RNA was subjected to reverse transcription, PCR amplified followed by Illumina index addition to create the final sequencing library.

### Data preprocessing

#### scID-seq

Sequence data from the NextSeq500 (Illumina) was de-multiplexed using bcl2fastq software (Illumina). Then, all reads were processed using our dedicated IDseq R-package ^20^. In short, the sequencing reads were split using a common “anchor sequence” identifying the position of the Unique Molecular Identifier (UMI) sequence, Barcode 1 (antibody specific) and Barcode 2 (well specific) sequence. After removing all duplicate reads, the number of UMI sequences were counted per barcode 1 and 2. Finally, barcode 1 (“antibody)” and barcode 2 (‘well’) sequences were matched to the corresponding antibody and well information.

Next, cell and antibody outliers were removed from the data. Specifically: cells with a total UMI-corrected count number of <25,000 or a median antibody count of <100 were removed. Cells were also removed if one or more antibodies had a UMI count >5 standard deviations above the population mean. Finally, antibodies with a median UMI count <40 were removed.

Antibody counts were normalized through TMM-normalization by, for each cell, scaling all counts with a normalization factor corrected library size. The normalization factors were calculated with the function *calcNormFactors* from the edgeR R-package ^47^ using the parameters sumTrim = 0.05 and logratioTrim = 0. Plate effects were corrected using the function *removeBatchEffect* from the limma R-package ^48^ using the plate number as batch indicator.

#### scRNA-seq

The sequence data were demultiplexed using bcl2fastq software (Illumina). RNA sequence data were processed using CELseq2 pipeline^49^ which includes demultiplexing based on cell barcodes, mapping to GRCh38 using Bowtie2^50^ and UMI counting by HTseq. The processed sequencing data were analyzed using the Seurat V4 package.

### Cell-state clustering

#### cell-state cluster assignment

To exclude treatment effects from affecting the clustering, clustering was performed only on so-called *cell-state markers*, i.e. (phospho-)proteins that are not significantly differentially abundant between treatments (Kruskal-Wallis test, comparing vehicle with p70S6Ki and p90RSKi, –log10 p-value <5) as previously described ^18^. Next, the TMM normalized data were z-score normalized on a per-(phospho-)protein basis to ensure equal contribution between (phospho-)proteins with high and low average number of counts. The z-score normalized data was transformed into an adjacency matrix using a shared nearest neighbor approach as implemented in the ‘RANN’ package ^51^, using k = 15.

Robust and reproducible “consensus” cell-state clusters were identified based on repeated application of the Leiden algorithm ^35^. The Leiden algorithm was run (with a resolution of 1.2) until it reached a stable configuration 1000 times. While the number of identified clusters varied from 7 to 12, approximately 70% of the runs resulted in nine clusters. To facilitate the determination of the consensus clusters, we filtered out all iterations of the Leiden algorithm where the number of clusters was different from nine. Next, a co-occurrence distance matrix of cells was calculated using the *cooccur* function from the ‘kmed’ package ^52^. Finally, hierarchical clustering on the co-occurrence matrix and cutting the resulting dendrogram with k=9 using the base R function *cuttree* resulted in nine consensus clusters.

To assess reproducibility of this pipeline, it was repeated 5 times. The vast majority of cells were consistently assigned to the same cluster in all 5 repeats. The 12 cells where this was not the case were marked as assigned to the cluster they occurred in most (in all cases 4/5 times), but marked as “ambiguous”.

#### cell-state cluster classifiers

Multinomial Logistic Regression (MLR), Random Forest (RF), and Support Vector Machine (SVM) classifiers were trained to assign cells to one of the 9 cell-state clusters. The classifiers were trained on the cell signaling markers, which were not considered in the cluster assignment. In addition, as a positive control models to predict the cell-states from the cell-state markers were trained. The classifiers used the z-score scaled protein counts of the TMM normalized data as input. The classifiers were evaluated using a nested cross-validation scheme with a five-fold inner and five-fold outer loop. This procedure was repeated five times. The Random Forest was optimized using randomly selected predictors (possible values 2, 5, 10), number of trees (possible values 500, 1000, 2000), and minimal node size (possible values 2, 5, 10) by exhaustively testing all parameter combinations in the nested cross-validation scheme. For the MLR model the penalty and model type were optimized and for the SVM the cost and sigma parameter. The MLR and SVM model parameters were tuned over a grid with 50 random parameter combinations.

#### Pathway activity score

To calculate the pathway activity scores, the TMM normalized data was rescaled to [0 – 10] so that each (phospho-)protein has the same weight when contributing to a particular score. To account for whether a factor/phosphorylation event activates or inhibits pathway activity, each (phospho-)protein was assigned to have a positive (+1), a negative (–1), or no (0) contribution to the score of each pathway (Table S5). The baseline score of each pathway was calculated by averaging the scaled antibody intensities of the (phospho-)proteins for all vehicle-treated cells in a cell-state according to the contribution of each (phospho-)protein to the pathway. These scores were subsequently scaled back to a ten-point scale to facilitate comparisons.

To calculate the fold-change in the pathway activity-score between baseline and inhibitor treatment, the pathway scores of individual p70S6Ki or p90RSKi treated cells were divided by the mean of the baseline score for each pathway-activity for each cell-state (cluster).

These fold changes with regards to the mean of the vehicle treatment of their cell-state were then used to compare the effects of the vehicle and inhibitor treatment with a Student’s t-test. For all tests, normality was assumed.

#### Identification of cell-state specific drug response

cell-state specific drug responses were identified by regressing the expression of each (phospho-)protein a in each cell *i*, *E*_i,a_, to the drug-treatment *T_i_* and cell-state *S_i_*, and include an interaction term, i.e. the following linear model was each (phospho-)protein using the base R function ‘lm’.

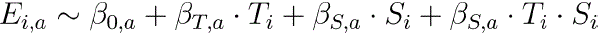

In the design matrix, vehicle treatment and cell-state 1 were used as reference states. Significance was assessed using a two-way ANOVA using the base R function ‘aov’. In addition, the significance of interactions for each cell-state-treatment combination individually was assessed using the p-values the interaction coefficients β_”,$_ was used. Multiple testing correction was performed using the Benjamini-Hochberg procedure as implemented in the base R function *p.adjust*.

#### Single cell Comparative Network Reconstruction

A full description of the single cell Comparative network reconstruction, including derivation of the underlying equations and validation on simulated and real data, can be found in our preprint on BioRxiv ^36^. Briefly, scCNR is a reformulation of our previous Comparative Network Reconstruction algorithm ^53^ that aims to quantify the interaction strengths 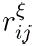 between the all protein pairs *i* and *j* in a signaling network, and how these differ between cell-states 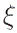. To this end, it exploits the stochastic variation in total and phospho-protein abundances. In a first order Taylor series, the deviation from the cell-state population mean of the activity and total abundance of protein *i* in cell *a*, 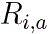 and 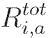 respectively, the sensitivity of the activity of protein *i* to changes in its total abundance, 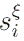, the effect of drug treatment *m*, 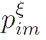, and the interaction strengths between proteins are related through the set of linear equation, one for each node in each cell:

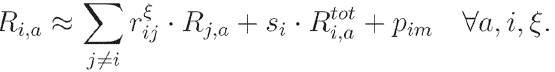

Here, the 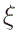 superscript refers to possible cell-state specific values of the parameters. scCNR solves an optimization problem to find a network (i.e. the values of 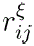) that fits these equations well while minimizing model complexity (i.e. the number of protein interaction in the network/nonzero values of 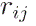) and the number of edges that differ in strength between the cell-states (i.e. have values for which 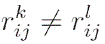 for different cell-states *k* and *l*). The strengths of the penalties on the number of edges in the network and the number of edges that differ in strength between cell-states is set by the hyperparameters *η* and *θ*, respectively. Prior information about the presence or absence of interactions can be easily incorporated as additional indicator constraints to the optimization problem, coded for by binary indicator variables *I_ij_*.

This optimization problem can be formulated as a Mixed Integer Quadratic Programming (MIQP) problem. The MIQP optimization problem is solved using IBM ILOG CPLEX solver (Version 22.1), which is freely available for academic use. Importantly, it guarantees optimal solutions within small numerical tolerances.

#### Identification of model topology

To reduce the search space, identification of network topology and interactions that differ between cell-states were decoupled. Initially, to obtain a suitable model topology, cell-state information was ignored, and the *R_i,a_* –values were calculated as the relative deviation in protein activity for the total population median. For nodes for which both the total and the phospho-protein abundances were quantified, the variation in total protein could be explicitly modeled as a perturbation to the node (the *R_i,a_*-term). For the nodes where we did not measure total protein abundance, we interpret this as an unobserved perturbation, resulting in worse ability to explain the variation in protein activity. To this end, the residuals corresponding to these nodes are down-weighted by a factor α, which is set automatically as described in ^36^. Drug treatments were modeled as negative perturbations to p90RSK (p90RSK inhibition) and RPS6 (p70S6K inhibition, as p70S6K is not measured itself).

A network of 65 canonical interactions and interactions obtained from PhosphositePlus ^37^ was included as prior information by adding indicator constraints of the form 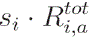, one for each edge in the starting network, to the optimization problem. Next, edges to be added to the network model were identified by solving optimization problem using *η* = 0.06 as this gave a good balance between model fit (as quantified by the RMSR and complexity, Figure S8b), ignoring cell-state information. This way, 34 edges are added to the model, resulting in a final model topology of 42 nodes and 99 edges.

The quality of the model topology obtained was assessed by comparing it to the fit obtained by 500 models with a random network topology. To this end, random models of the same complexity (i.e. 99 edges) were created by randomly setting 99 indicators *I_ij_* = 1 and all others indicators to 0. By solving the MIQP optimization problem under these constraints, the interaction strengths and thus also RMSRs of these models were obtained. These could than be compared to the RMSR from the actual model (Figure 4b)

#### Identification of cell-state specific edge strengths

To identify which edges differ between the cell-states, scCNR with cell-state information included was run. To this end, the input values of *R_i,a_* and *R_i,a_* were calculated as the relative deviation in protein activity from the cell-state median. The network topology as obtained above was provided by setting the 99 indicators corresponding to included edges to 1 and all other indicators to 0. The hyperparameter penalizing differences between cell-states, $\theta$, was set to 0.2 as this gave a good balance between the number of differences between the networks and the reduction in RMSR (Figure S8c). Otherwise, the optimization was run as described above. This resulted in a network with 34 differences between the networks: 31 of the differences were edges, 2 differences in sensitivity to the total protein abundance, and 1 a difference in the direct effect of a drug treatment on its target.

The bootstrap analysis to assess the robustness of differing edge detection (Figure S8d) was performed by randomly sampling 546 cells (i.e. the original number of cells) *with* replacement and rerunning the MIQP optimization again with *θ* = 0.02 (i.e. the original value). This procedure was repeated 100 times. From this, the fraction of bootstraps in which an edge is predicted to be cell-state specific can be obtained.

The permutation analysis to assess the statistical significance of edge strengths deviations (Figure 4d,e and Table S4) was performed by randomly shuffling cell-state labels. To ensure identical distribution of cell-states across treatments, this was done for each treatment separately. Next, the cell-state median and R*_ij_* values were recalculated. The model topology and the edges that differ were all fixed by constraining all indicator variables to be the value they had in the original model. Subsequently, the MIQP optimization was run to obtain the edge strengths for the permuted data. This was repeated 1000 times to obtain a null-distribution of edge-strength-deviation from population mean.

#### Enrichment analyses

For the enrichment analyses presented in Figures 4h and S8e, we considered all nodes with a significant interaction term according to the ANOVA analysis (Table S3) and edges that are identified as having cell-state specific interaction strengths in the general scCNR optimization, to obtain the contingency tables. Odds ratios and p-values were calculated using one-sided Fisher Exact tests.

For the cell-state resolved enrichment analysis, we considered all node-cell-state pairs that have a significant coefficient for their interaction term in the regression analysis (using cell-state 1 as reference), and all edge-cell-state pairs where the deviation in interaction strength significantly differed from the null-distribution obtain from the cell-state label permutation analysis. This way, we obtained the contingency for edge and node enrichment, respectively. Odds ratios and p-values were calculated using one-sided Fisher Exact tests.

## Data and code availability

Sequencing data for the scID-seq and scRNA-seq experiments produced for this work are deposited in the Gene Expression Omnibus (GSExxxx). Moreover, the TMM normalized data are available as supplementary data associated with this manuscript. Data analysis code is available on https://github.com/jessievb/IDseq and https://github.com/evertbosdriesz/scIDseq-CNR

## Figures and Legends

**Figure S1.**
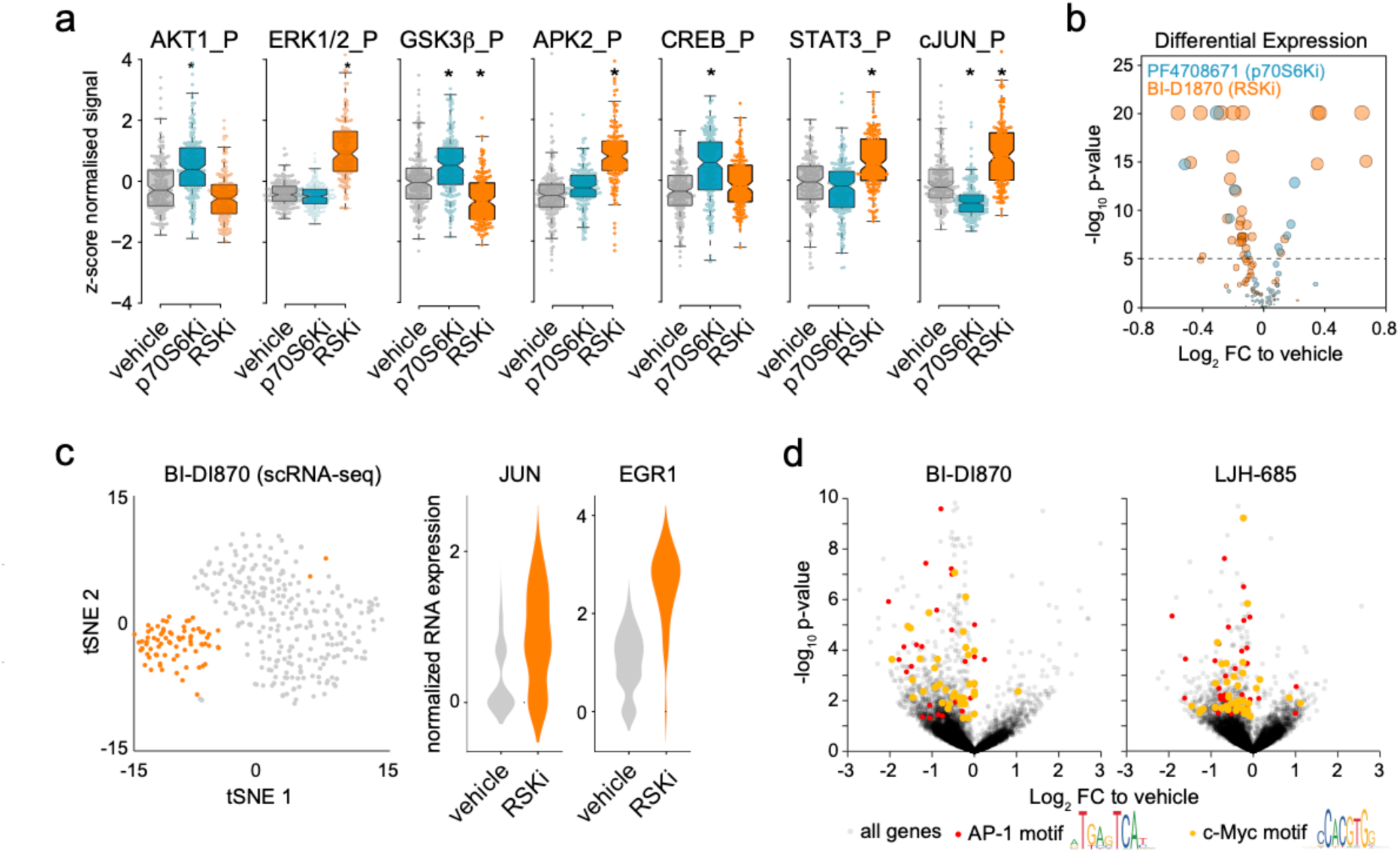
Single cell profiling of signaling protein activity to targeted inhibition. **a.** Boxplots of phospho-protein abundance of core EGFR pathway components upon drug treatment. **b.** Volcano-plot the –log_10_ p-value and –log_2_ fold change of (phospho)-protein abundance in response to targeted, compared to vehicle control. **c.** tSNE representation of scRNA-seq profiling (left) and a violin plot of the JUN and EGR1 normalised expression (right) of human keratinocytes treated with vehicle, or BI-D1870 (p90RSK inhibitor). **d.** Volcano plots of differential expression based on scRNA-seq data of human keratinocytes treated with vehicle or the p90RSK inhibitors BI-D1870 or LJH-685. Genes with known AP-1 or c-Myc motifs in their promoter region are indicated as red and orange data points, respectively.

**Figure S2.**
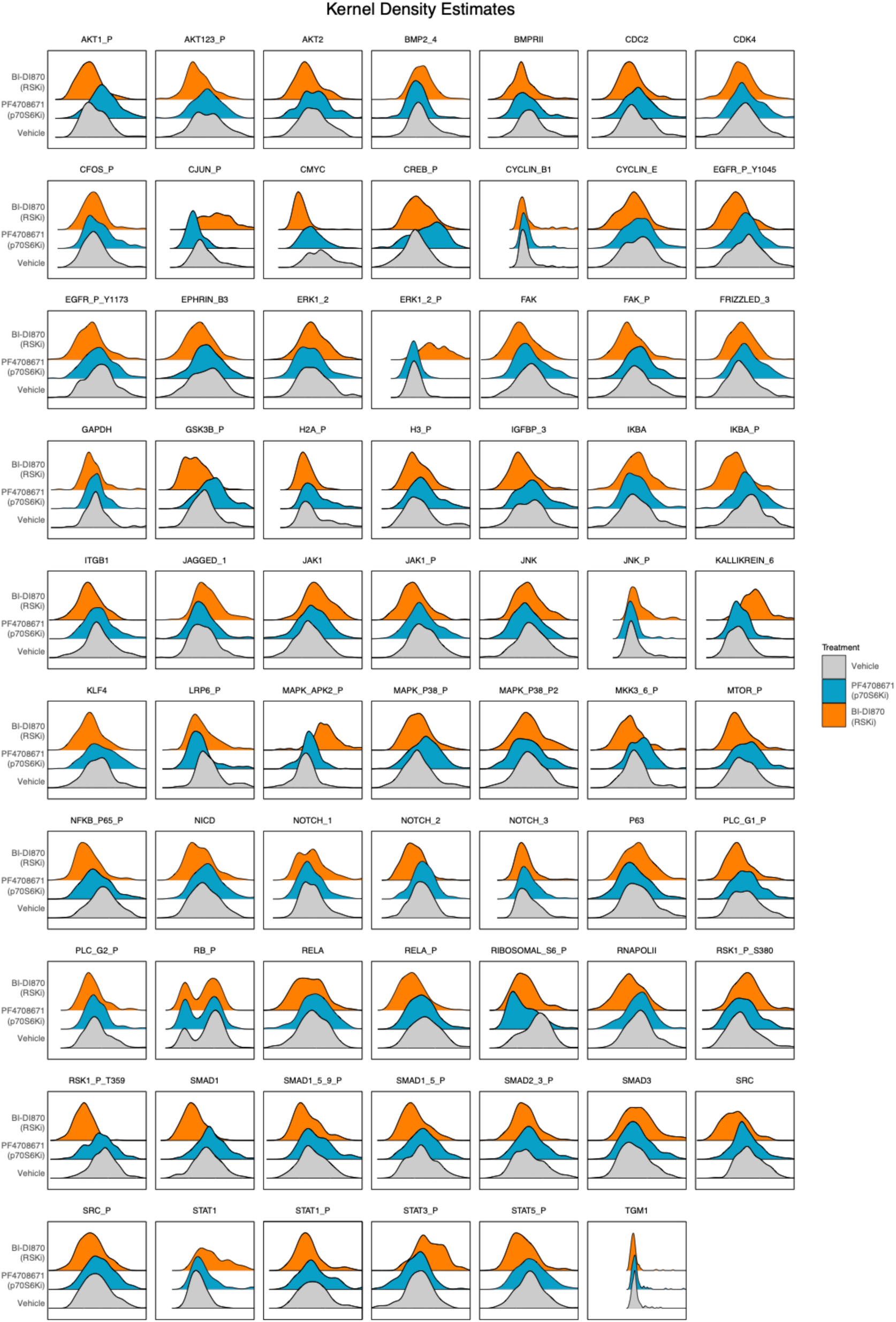
Distributions of (phospho)-protein abundances. Ridgeline plots of the abundances of all (phospho-)protein abundances in the panel, separated by treatment.

**Figure S3.**
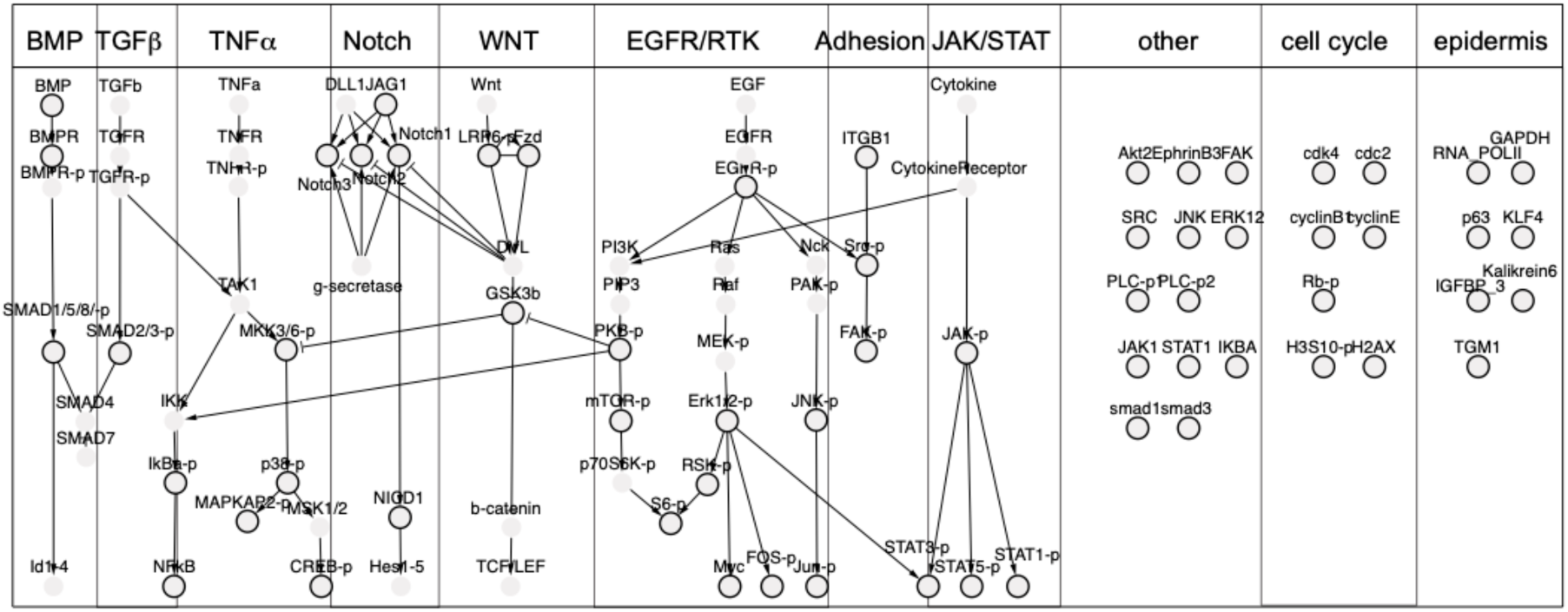
Node identities of the biochemical signaling network on which the scID-seq data is projected. Edges between Nodes ((phospho-)proteins) are drawn based on literature-established interactions.

**Figure S4.**
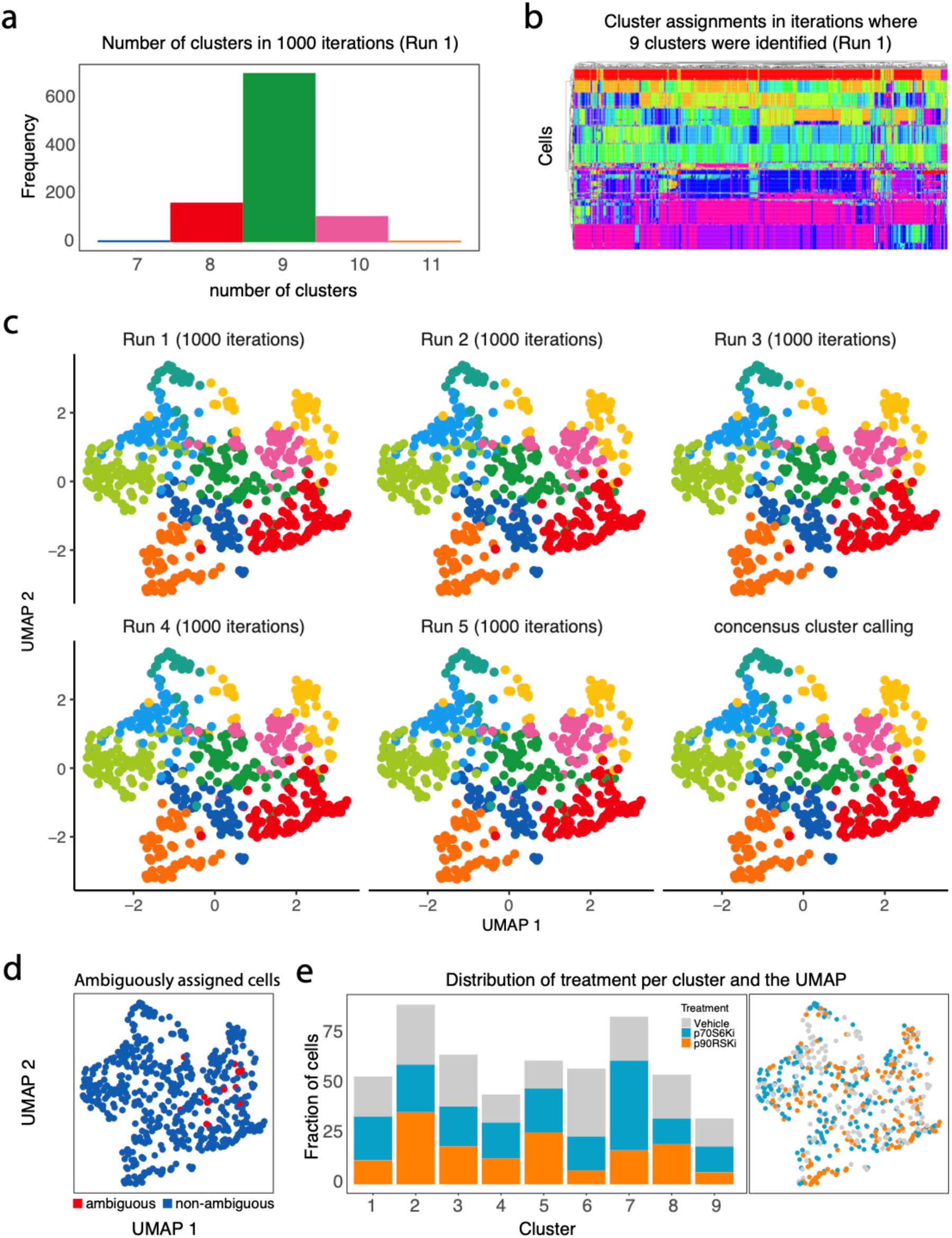
Robust clustering of cells into distinct cell-states. **a.** Distribution of the number of clusters identified by the Leiden clustering algorithm with 1000 random initializations. **b.** Hierarchical clustering on the co-occurrence matrix of cells obtained by repeating the Leiden clustering algorithm 1000 times. Instances in which 9 clusters were obtained (>60% of iterations) were included in the co-occurrence matrix. Cells were clustered into groups by cutting the dendrogram at k=9. This procedure was repeated 5 times. **c.** UMAP projection of the cells color coded according to their cluster assignment in each of the 5 runs, and the resulting consensus cluster calling. Only the cell-state markers were used in determining the UMAP coordinates. **d.** UMAP projection of the cells color coded by whether the cells were assigned to the same cluster in each of the 5 runs (blue points), or not (red points). **e.** Left panel: Barchart indicating the fraction of cells in each cluster, color coded by treatment. Right panel: UMAP projection of the cells color coded by treatment. Together, this shows that cells from the different treatments are fairly randomly distributed over the clusters, and also not clearly separated in the UMAP. This is what we would expect, since the clustering is based on markers that do not show a response to the treatments.

**Figure S5.**
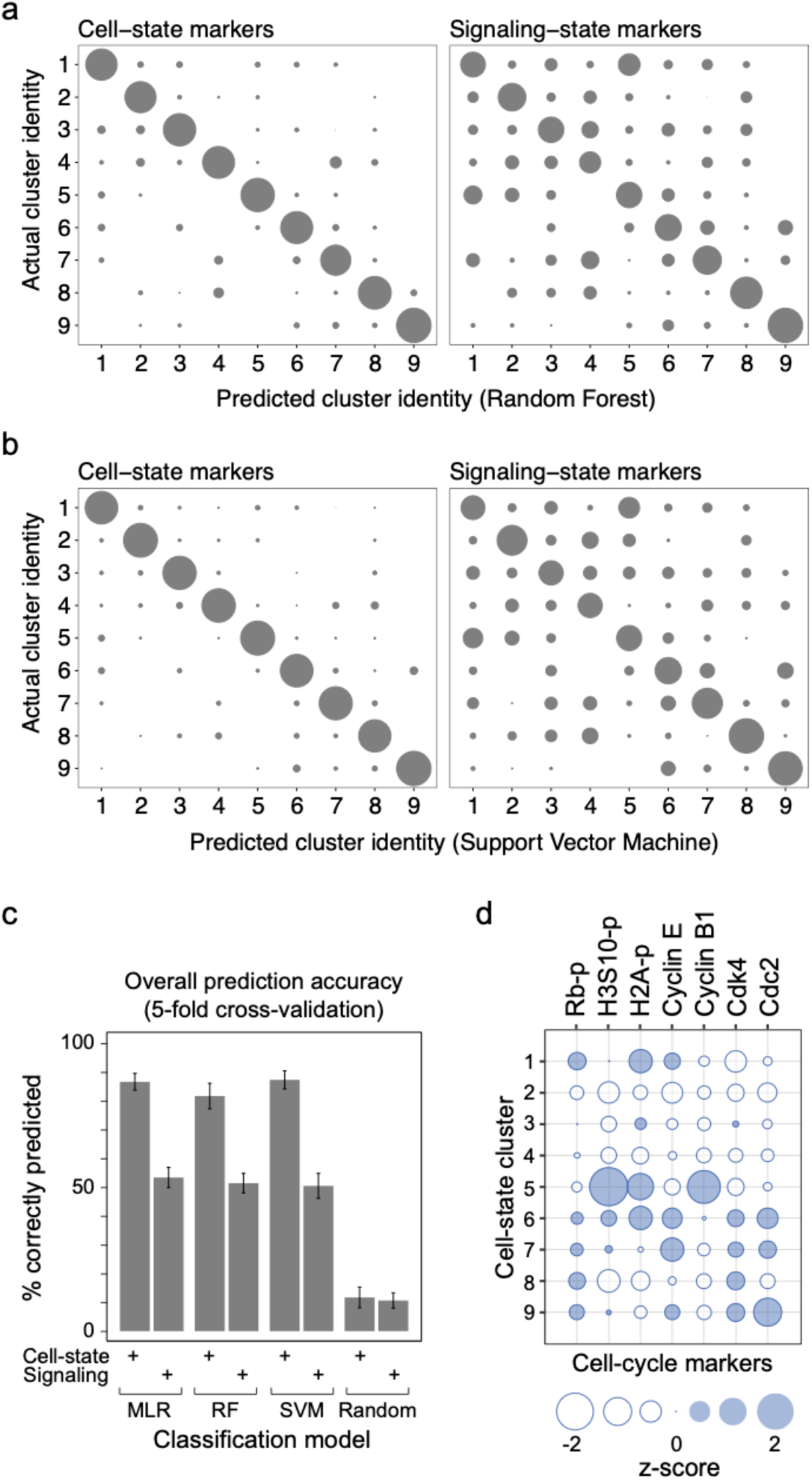
Signaling proteins not used in cell-state assignment nonetheless show cell-state dependent behavior. a,b). Bubble chart comparing the cluster identity as predicted by a Random Forest (RF, panel **a**) or Support Vector Machine (SVM, panel **b**) model using the cell-state marker (left panels) or signaling-state markers (right panels) with the actual cluster identity. Model performance was assessed using a nested cross-validation scheme of 5 inner and 5 outer folds, which was repeated 5 times. **c.** Bar charts comparing the prediction accuracy of MLR, RF, or SVG with those of a random model. Error bars represent standard deviation. **d.** Bubble chart representing relative levels of the indicated cell-cycle markers. Bubble size indicates average z-score of the cells belonging to the indicated cell-state in the vehicle condition, negative scores are indicated by empty bubbles.

**Figure S6.**
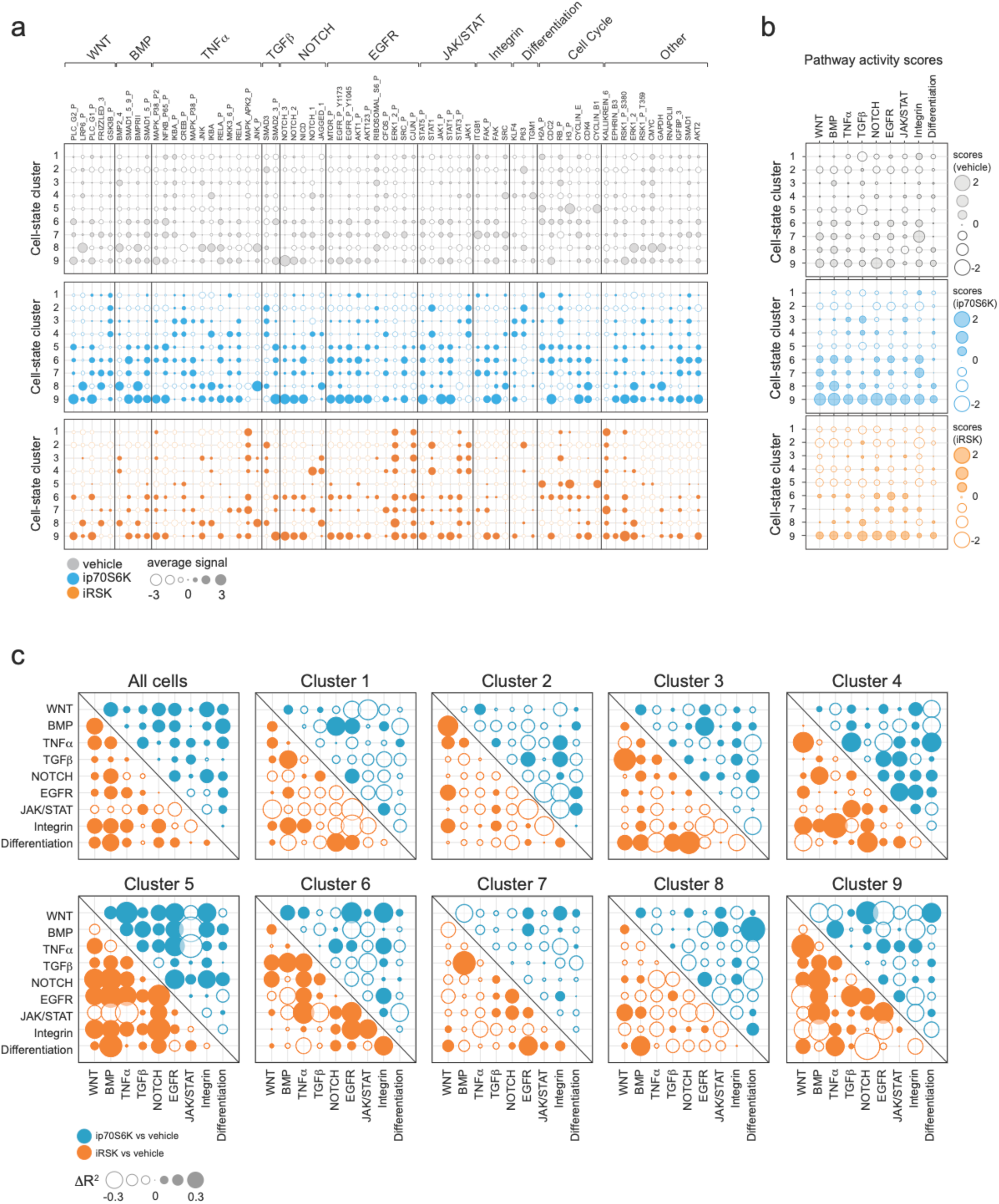
The response to targeted inhibition is cell-state dependent for many (phospho)-proteins. **a.** Bubble chart indicating the abundance of all (phospho-)proteins per cell-state and treatment. Bubble size indicates average z-score normalized abundance of all cells belonging to the corresponding treatment and cell-state. Colors correspond to treatments. Positive z-scores are indicated by filled bubbles, negative z-scores by empty bubbles. **b**. Bubble chart indicating the Pathway activity scores per treatment and cell-state. Bubble size indicates average activity score of all cells belonging to the corresponding treatment and cell-state, and negative scores are indicated by empty bubbles. **c.** Bubble chart indicating the difference in variation explained (linear regression, R^2^) between vehicle and treated conditions per pathway, per cell-state cluster. Bubble size indicates the difference in R^2^, negative values are indicated by empty bubbles.

**Figure S7.**
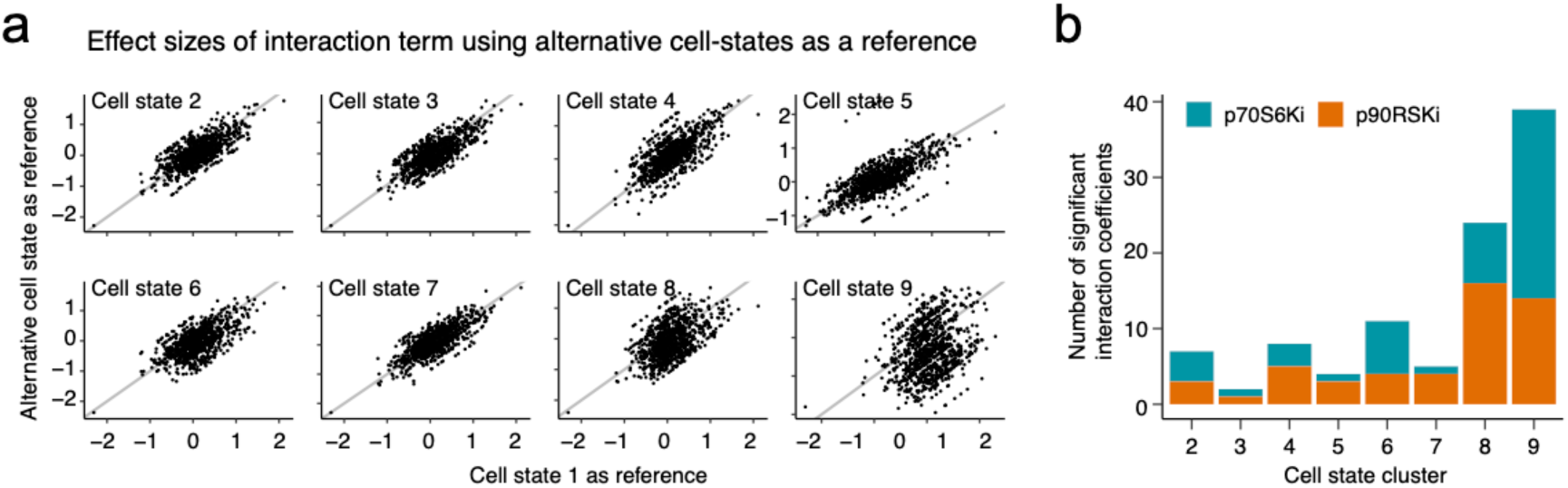
Cell-state dependent drug response analysis. **a.** Scatterplots of comparing the effect size of the drug treatment-cell-state interaction term using different reference cell-states in the linear model. Each panel compares a linear model with a different reference cell-state with one using cell-state 1 as reference. Each point corresponds to a cells-state-drug treatment combination. **b**. Bar chart indicating the number of significant (p<0.05) interaction terms per cell-state, colors represent the different treatments.

**Figure S8.**
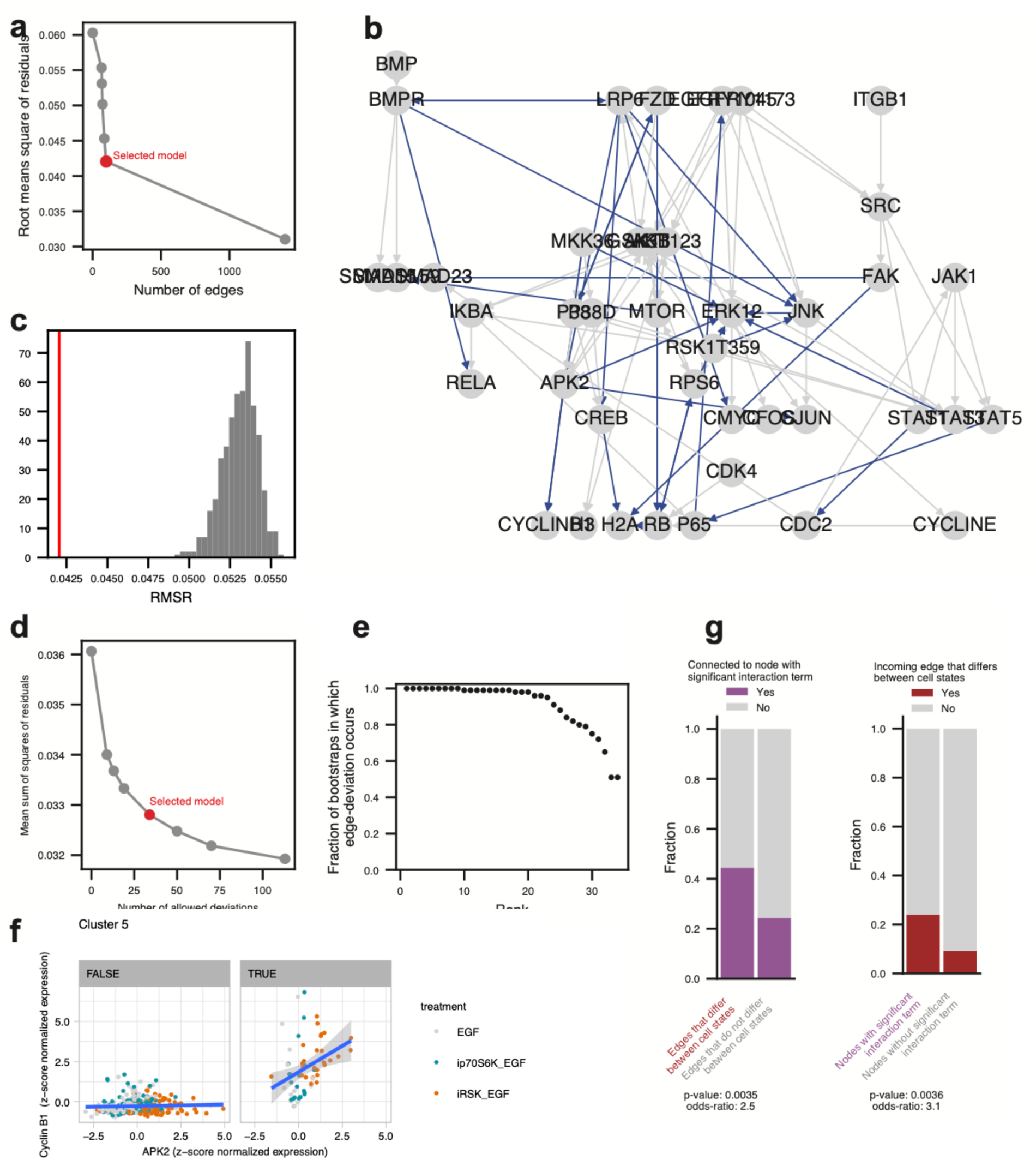
Single cell Comparative Network Reconstruction. **a.** The Root Mean Square of Residuals (RMSR) of the model fit obtained by scMRA MIQP optimization as a function of the number of edges added to the prior network. The number of edges added to the prior model was varied by varying the hyperparameter *η*, which sets the penalty on the number of edges in the network. The red point indicates the model that we selected. **b.** Topology of the selected model. Gray edges indicate interactions based on prior knowledge, blue edges indicate interactions that were added in the scMRA optimization. **c.** Distribution of the RMSR of models optimized using a random topology (gray histogram) and with the actual model as depicted in panel b (red vertical line). **d.** RMSR of the scCNR model fit, as a function of the number of interactions that are different between the 9 cell-states. The number of differences was varied by varying the hyperparameter *θ*, which sets the penalty on the number of edges that are different in strength between the cell-states. The red point indicates the selected model. **e.** Fraction of bootstraps in which an edge that was identified to have different interaction strengths between cell-states. Each point represents an edge that was identified to have different interaction strengths in the actual model (i.e. the red edges in Figure 4a). **f.** Scatterplot of Cyclin B1 expression against phospho-APK2 in Cell state cluster 5 (right panel) or in all other cell states (left panel). **g.** Visual representation of the contingency tables describing the connection between cell-state dependent drug response and interaction strengths. Left panel: The bars indicate the fractions of the node-cell-state pairs that had a significant interaction term in the linear regression model, separated by whether or not they have an income edge that is significantly different from the population mean for that cell-state, as assessed by the permutation analysis. Right panel: Fraction of edge-cell-state pairs where the edge strength is significantly different from the population mean for that cell-state, separated by whether or not they connect (upstream or downstream) to a node showing that has a significant interaction term for that particular cell-state.

